# *IQC:* A Novel Criterion for Assessing Feature Selection Stability in High-Dimensional Analyses

**DOI:** 10.64898/2026.09.26.754480

**Authors:** Tristan A. Moxley, Benjamin J. Ridenhour

## Abstract

In recent years, the issue of reproducibility in scientific studies has been at the forefront of scientific discussion. Many of these issues arise due to poor employment of statistical methods, making it crucial to address these knowledge gaps for frequently used methods. Elastic net regression (ENET) is one such algorithm used frequently in biology and public health due to its ability to select important predictors (also known as “features”) from large data sets. However, ENET struggles with consistent feature selection and hyperparameter choice in these applications due to low observation counts and noisy data. This limitation hinders ENET’s effectiveness in biological research due to low model reproducibility and predictor interpretability. We propose the Instability Quotient Criterion (IQC) as a metric to quantify the consistency of selected features for feature selection models such as ENET, with a specific emphasis on biological applications. IQC leverages ensemble modeling and Shannon’s entropy in order to determine the stability of selected features derived from ENET models. We demonstrate IQC’s robustness via simulation studies, showing that IQC is capable of detecting significant decreases in model stability with lower observation counts, higher dimensionality, and higher proportions of noisy signal. We additionally utilize simulation to provide a naive threshold for low versus high stability. Finally, we applied IQC to a case study to illustrate how our method can be integrated into a larger analysis pipeline when looking at microarray gene expression data. IQC indicated stability issues for differentially-expressed genes when classifying acute leukemia subtypes in spite of high classification success. Lastly, we discuss how model instability impedes interpretation of biologically significant genes for discerning cancer subtypes. Overall, we present IQC as a robust evaluation metric that can easily be incorporated into high-dimensional biological analyses.

**Author summary:** Addressing the reproducibility crisis is paramount for bolstering confidence in scientific studies. In biological settings specifically, making consistent predictions when using machine learning techniques is vital. Methods such as elastic net regression (ENET) often suffer from inconsistent predictor selection when applied to biological data due to high dimensionality, noise, and low observation counts. Thus, the ability to detect when predictor selection is unstable is valuable for researchers in biology and health. We present the Instability Quotient Criterion (IQC) as a novel criterion to assess model stability for elastic net regression. This criterion gives a value between 0 and 1 that displays how consistently the same predictors are chosen. We find through simulation studies that IQC is robust in its detection of a stable versus unstable model when considering its selection of important predictors. We also demonstrate IQC’s usefulness in a larger modeling pipeline when applied to gene expression data. Our findings show that IQC is a valuable tool for inferring the robustness of biological predictors with more nuance and precision.

## Introduction

In recent years, the issue of reproducibility in scientific studies, including ones pertaining to biological discovery and machine learning, has been at the forefront of scientific discussion [1]. Many reproducibility issues arise due to insufficient computational training and improper employment of statistical techniques, both of which can result in the acceptance of uncertain results with overconfidence [2–4]. Addressing this aspect of the reproducibility crisis involves bolstering knowledge of the limitations of statistical methods, especially ones used frequently in biological research.

One such limitation that results in reproducibility issues is model instability, especially when using sophisticated machine learning techniques. “Model stability” refers to the capacity of a statistical model to make consistent predictions, select the same important predictors, and/or produce the same outcomes [5]. Stability is often compromised due to low observation counts, high data dimensionality, or incorrect model applications [2, 6, 7]. Previous research cites the importance of checking stability to aid in reproducibility for these analyses, especially when applied to biological research and health outcomes [1, 5]. As such, stability is a primary focus for addressing gaps in knowledge with respect to machine learning and its usage in biological research.

Elastic net regression (ENET) is a widely-used machine learning algorithm that utilizes penalized least squares regularization for selecting important predictors (also known as “features”) from large data sets [8]. This method has been applied to a wide array of biological and health-oriented studies, such as predicting long COVID-19 and metabolites responsible for oral health in children [9–12]. ENET is especially useful for data with high-dimensionality (a common trait of many biological data sets) due to its ability to both a) set variable coefficients to 0 via lasso penalization (*l*_1_) and b) shrink collinear variable magnitudes towards each other (known as the “grouping effect”) via ridge penalization (*l*_2_), resulting in more robust model performance compared to each penalty alone [8].

Though not as cutting-edge as current advancements in deep learning, ENET remains an advantageous modeling infrastructure in biological research due to its interpretability and background as a method designed for biological data. Deep learning methods, while highly precise, often struggle with interpretability, being touted as “black boxes” [5, 13]. Conversely, methods such as ENET have much more interpretable results given their foundation in regression and statistical theory. Additionally, ENET was principally developed for usage in high-dimensional biological analyses; the foundational literature that established and first-utilized ENET (along with its component methods lasso and ridge) specifically addressed improving modeling when applied to genome wide association studies (GWAS), cancer subtype classification, and ecological modeling [8, 14–18]. Further, ENET was explicitly developed to solve lasso’s random selection of a single variable from a collinear group [8]. Thus, ENET remains an ideal modeling choice when applied to high-dimensional biological data.

Despite these benefits, ENET often struggles with consistent feature selection when data are noisy [6, 7] and when observation counts are low [19]. Both of these issues are often impossible to avoid during data collection. For example, infectious disease data are prone to data reporting errors from poor reporting and (in some cases) asymptomatic hosts [20]. In bioinformatic settings, DNA sequencing via 16S amplicon or shotgun methods often present unavoidable contamination during the extraction process [21]. And, downstream genomic analyses for microbial abundance and gene expression, for instance, can rarely obtain sufficient sample sizes to perform robust modeling given the large number of species or genera present in these data [22–24]. Although ENET is designed for such feature selection in cases where *p >> n* (i.e., a data set with many more predictors than observations), high error/variance and can significantly diminish ENET’s predictive capabilities [25, 26]. Reduced predictive capability persists even if many of the noisy predictors are independent of the response variable [27] and is often exacerbated by low observation counts inducing high variance within each feature [7, 28].

ENET uses cross-validation to prevent the model from overfitting the training data; however, this can often induce further model instability. Data splitting is a common form of cross validation, where the model is “trained” using a majority of the data set, then validated on the remaining data, to ensure the model generalizes well [29]. Several studies aim to determine an optimal ratio for training and testing data splits; however, standard practice opts instead for simple ratios, such as 70:30 or 80:20 [29–31]. Though this step is intended to improve the final model, cross validation can fail to generalize a data-poor model [19], as low observation counts and/or high dimensionality induces high variance in the training data. This higher variance results in selected features that vary wildly, thus making stability and interpretation unreliable [32].

Improving reproducibility, in part, involves a better understanding of statistical methodology and better data management practices [33]. When specifically focusing on methods such as ENET, this can be approached via the development of quantifiable metrics to assess how reliably features are selected. Much like with existing measures of model accuracy such as mean-squared error, a univariate metric provides an easily-interpretable way to assess feature stability, which can thus aid in improving reproducibility for more sophisticated statistical procedures. When considering ENET, previous studies have employed stability selection as a means to make model output more reproducible [34], and several discipline-specific techniques have been proposed to improve ENET’s modeling infrastructure [7]. Additionally, more broad applications of stability analyses have been proposed for health-oriented analyses [5]. However, development of a diagnostic measure to understand the consistency of model output has not been explored for ENET [3].

We propose a novel metric of stability called the Instability Quotient Criterion (IQC) which assesses the consistency of selected features in elastic net regression models. IQC relies on consensus modeling via ensemble techniques, as well as evaluation via Shannon’s entropy [35], to assess feature selection reliability. We developed and validated this method using a robust data simulation algorithm which tailors the number of predictive features, proportions of true signal to noise, and number of observations. We analyzed three separate data configurations based on the ratio of signal-to-noise, while computing IQC for decreasing observation counts. Finally, we applied our method to leukemia microarray data from the original ENET publication to analyze the stability of selected genes in predicting leukemia subtype [8, 36].

## Results

A brief overview of the IQC procedure will be described here, with the full derivation being available in the Materials and Methods section. IQC is an algorithm that computes stability via ensemble modeling and Shannon’s entropy [35]. We begin by permuting the original data *X* into unique training and testing sets *i* times (for our purposes, we generate *i* = 100 permutations). For each permutation, we perform ENET using each permutation’s respective training set, then validate on its respective testing set. We label these as submodels for IQC, with each individual model being referred to as an iteration (or permutation). Each of these submodels possess unique final predictor sets, model accuracy, and model hyperparameters. Once all 100 submodels are generated, a “reference” model is computed as an aggregation of all model iterations. We then compare each model iteration to the reference model, noting which features match the reference with respect to magnitude (i.e., zero versus non-zero coefficient) and sign. Finally, IQC is computed as:

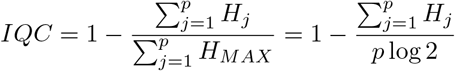

where *p* is the number of features, *H* is the entropy for all *p* predictors according to how consistently they align with the reference model, and *H*_*MAX*_ is a theoretical “maximum-entropy” model where each inclusion probability is equal to 0.5 (i.e., each feature is completely randomly retained or removed from the model iteration).

Using generative models, we simulated responses for all *p* predictors (which comprises all true signals and noise) via ENET with Gaussian errors. To test IQC’s ability to act as a diagnostic metric, we first began by manipulating the signal-to-noise ratio and numbers of observations in tandem. We observe the behavior of IQC for four different total feature space sizes: *p* = 60, *p* = 100, *p* = 140, and *p* = 200. To assess the impact of sample size on IQC, we varied sample size from 30 to 200 in steps of 10 observations. IQC is subsequently calculated at each step. To account for the simultaneous effects of feature dimensionality and observation counts, we report all IQC values with respect to the ratio of observations to feature (OTF) counts. We define OTF as 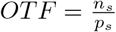 (where *n*_*s*_ and *p*_*s*_ are the number of observations and features for a given simulation *s*, respectively).

Fig 1 displays the behavior of IQC when the OTF ratio is incrementally increasing for three different proportions of true features to noisy signal; S1 Table shows the full set of IQC values for each simulation configuration. Across each signal-to-noise simulation, we observed a consistent increase in IQC as sample size increases. We also observe that generative models with higher complexity (i.e., models with more features) tend to have lower IQC values overall compared to generative models with smaller feature spaces. Interestingly, reporting each simulation with respect to the OTF ratio as shown in Fig 1 shows that IQC exhibits very consistent behavior as OTF increases. From a ratio of 0.5 to just under 3.5, IQC steadily increases, and each simulation line overlays each other with minimal variation. Slight differences do occur for differing signal-to-noise ratios, and will be discussed individually.

**Fig 1.**
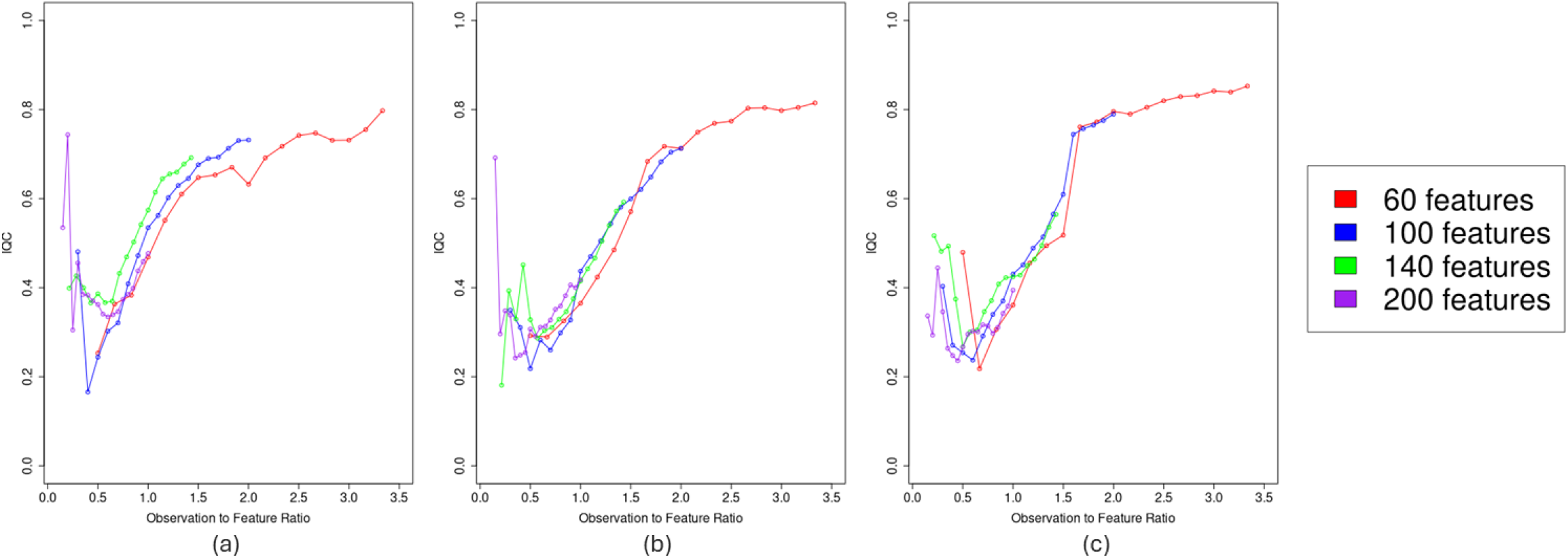
Behavior of IQC across each simulation design. (a) 25% Signal-to-Noise ratio, (b) 50% Signal-to-Noise Ratio, (c) 75% Signal-to-Noise Ratio. IQC values are computed for four configurations of the simulated data (200 observations 60 features, 200 ×100, ×200 ×140, and 200 ×200), with three separate proportions of true signal to noise (25%, 50%, and 75%). Each point on the line represents a unique IQC value when varying the number of observations per data configuration. The x-axis is rescaled to be the Observation to Feature Ratio at each step (defined 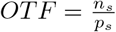, where *n*_*s*_, *p*_*s*_ are the number of observations and features for a given simulation *s*, respectively) to control for differing ratios of observation counts and features per configuration.

We also assessed the accuracy of each data configuration with respect to how close each reference model aligned with our true predictor set used in the generative model. Fig 2 displays receiver-operating characteristic (ROC) curves for each of the four feature space configurations, across each of our simulations. Given the thresholds for each ROC curve are empirically derived from our simulations, we utilized LOESS regression (local polynomial regression fitting) to approximate the shape of each ROC curve [37]. Despite having high variability in true positive rates (TPR) when sorting false positive rates (FPR) in ascending order, each ROC curve lies above the random classifier center line (as indicated by the black dashed lines in each subplot), indicating generally positive classification (i.e., elastic-net properly chooses true predictors). Additionally, several of the ROC curves are truncated prior to reaching an FPR of 1, meaning that we observe a lower incidence of false positives compared to true positives. Overall, while the individual ROC points show variability due to being derived from empirical data, our simulations display sufficient accuracy relative to the true features and noisy predictors of our simulated data sets.

**Fig 2.**
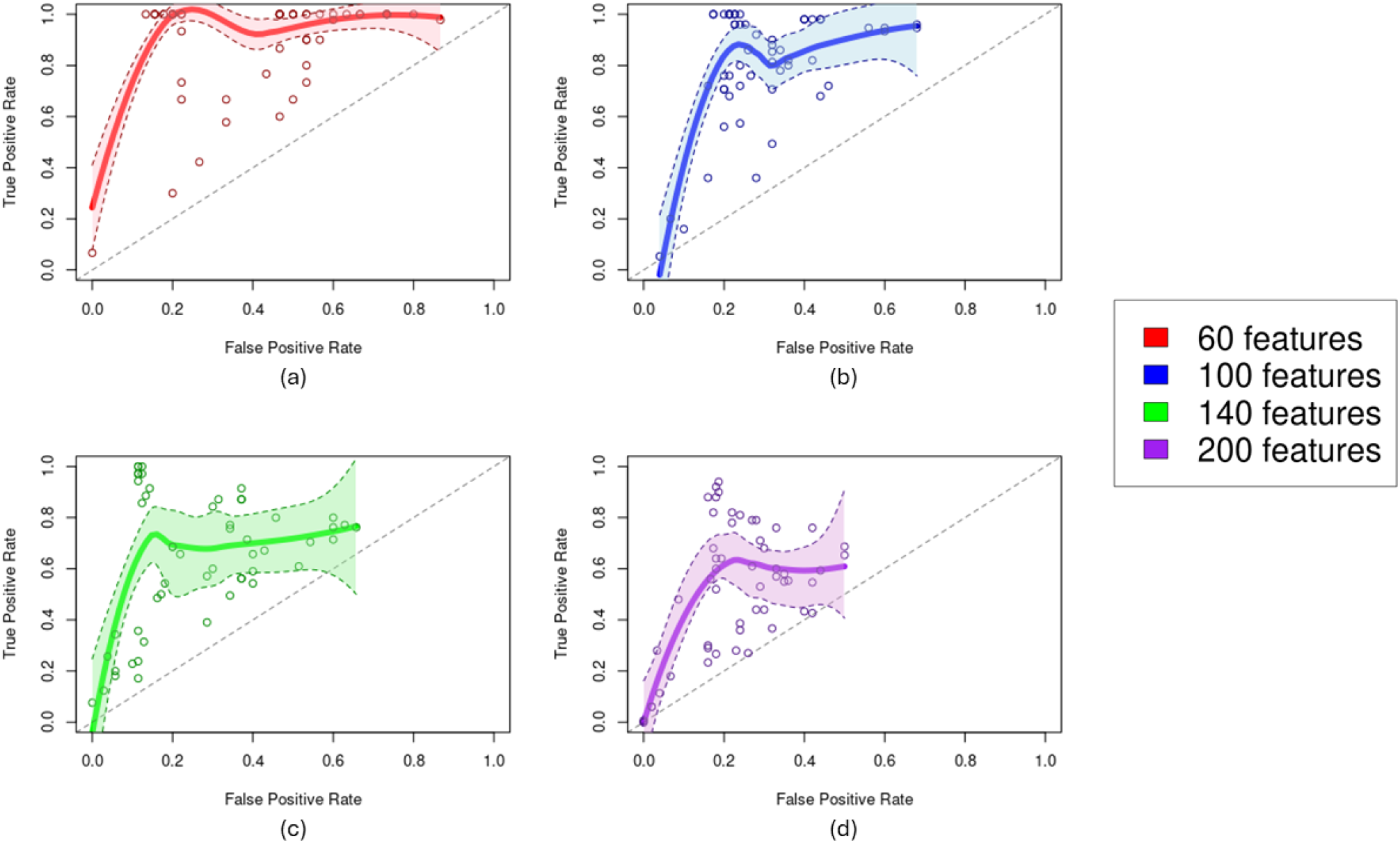
Fitted receiver-operating characteristic (ROC) curves to assess the accuracy of each IQC model iteration when compared to the true simulated model. (a) 60 features, (b) 100 features, (c) 140 features, (d) 200 features. ROC curves are computed for each feature space by aggregating the confusion matrices for all OTF and signal-to-noise ratios. True and false positive rates are the proportion of model iterations per data configuration that retain true features and remove noisy features, respectively.

For a 25% signal-to-noise ratio, we find that the IQC for these models increases suddenly for especially low OTF ratios. This effect is magnified as the total number of features present in the model increases. For instance, the blue line in Fig 1a, representing *p* = 100, steadily decreases in IQC value up until an OTF ratio of approximately 0.5, at which point its IQC value increases suddenly. Similarly, the purple line, representing *p* = 200, appears to have steadily increasing IQC values when the OTF ratio gets lower than 0.5, then suddenly spiking at a ratio of approximately 0.4. This effect appears to display higher “stability” for the higher-dimensional models when OTF ratios are especially low according to IQC alone. When observing the accuracy plots, we note that models with fewer total features had much higher accuracy when decreasing the OTF ratio.

For 50% signal-to-noise, we observe similar patterns with respect to the dimensionality of the data sets used. The models with more overall features tended to have lower IQC values compared to models with less features. For instance, the red line representing *p* = 60 peaks at an IQC value of around 0.8, and generally has the highest overall IQC until achieving an OTF ratio of approximately 1.75. Conversely, the green line representing *p* = 140 begins at an IQC of around 0.55, and is consistently more unstable according to IQC until the OTF ratio becomes very low.

Finally, 75% signal-to-noise displays the most regularity of the three data configurations. We see similar stratification with respect to the total number of features per model, with lower-dimensional models exhibiting better stability than higher-dimensional models. Unlike the other two plots, however, this configuration does not have the same extreme peaking in IQC when the OTF ratio becomes very low. Interestingly, we see a distinct corner where IQC begins to decline at approximately an OTF ratio of 1.5, whereas each other configuration exhibited a generally smoother decrease. This lack of peaking at low OTF ratios is possibly caused by the higher degree of true features present within the data.

We visualize the simulated distribution of IQC values across all data configurations in Fig 3. Theoretically, IQC can take on a value between 0 and 1; in practice, we observed a minimum value of 0.1657, and a maximum of 0.8526. The median IQC value was 0.4403, with the mean value being 0.4847. According to the histogram, the distribution has a slight right skew and is bimodal. Given these values, and the distribution of IQC, we recommend the following thresholds for low, moderate, and high stability: For low stability, we define the threshold as being below 0.35. This is indicative of all values at or below the first mode of the distribution. For moderate stability, we define this as being any value between 0.35 and 0.65, representing all values between the first mode and quartile 3. Finally, all IQC values above 0.65 are considered highly stable.

**Fig 3.**
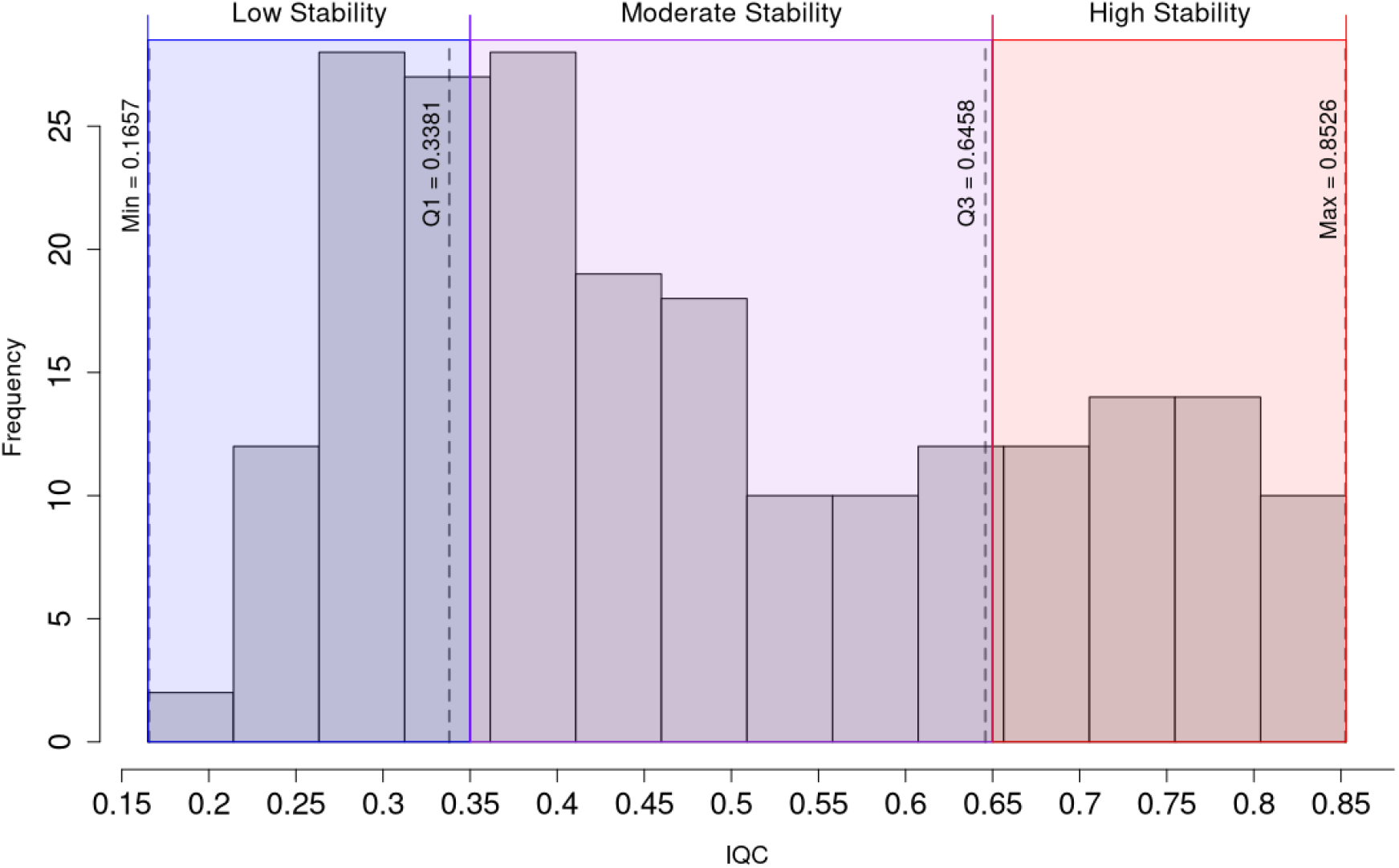
Distribution of IQC values derived from simulation study, along with proposed thresholds for low, moderate, and high stability. The blue region indicates low stability (IQC between 0 and 0.35), the purple region indicates moderate stability (IQC between 0.35 and 0.65), and the red region indicates high stability (IQC between 0.65 and 1). Values for the quartiles (minus the median) are overlaid on top of the histogram to highlight their role in defining IQC thresholds, specifically Q1 and Q3.

### Case Study: Classifying Leukemia Subtypes via Microarray Gene Expression

Similar to [8], we tested the IQC algorithm against microarray gene expression data gathered by [36]. These data contain the normalized relative expression of 7129 genes across 72 total individuals, with each individual being labeled as either having Type 1 (acute lymphoblastic leukemia, or ALL) and Type 2 (acute myeloid leukemia, or AML) leukemia. The response vector *Y* corresponds to *I*(*ŷ >* 0.5), where *I*(·) is the indicator function; this transformation codes our fitted values *ŷ* as AML if the value is greater than 0.5, and ALL otherwise. To provide a similar comparison to [8] and [36], we tested the IQC algorithm on a pre-filtered subset of the original data (*p* = 1372 genes). The approach to filtering is described further in the Materials and Methods section.

Table 1 displays the result of the IQC algorithm on 100 iterations of the ENET model for these data. Given the usage of logistic ENET, we based the decision rule for each IQC submodel on classification error as opposed to MSE (see Methods section). We found that no individual model iteration had a testing set classification error higher than 22.7% (with an average classification error of less than 3% for both the Full and Filtered Models), meaning that each model performed well at correctly classifying patients into their true leukemia subtype. However, the overall stability according to IQC is just within the lower bound of “moderate” (according to our proposed thresholds) at 0.386 for the full set of genes, and slightly higher at 0.390 for the filtered data. This implies that, while each model iteration accurately identifies proper leukemia subtypes, it does so while selecting different genes as the “significant” predictors with considerable frequency. We visualize this instability in Fig 4 by comparing the distribution of retained genes and the classification accuracy for each ENET submodel. While many of the model iterations preserve only 0 to 50 genes (as indicated by the tallest bin of the histogram), 60% of all model iterations preserve between 50 and 450 genes, and roughly 10% preserve all 1372. This further illustrates that each ENET submodel, while performing the exact same computations, is selecting completely different sets of predictors for classifying leukemia subtype. Further, Fig 4b shows nearly every ENET submodel predicts leukemia subtype with at least 95% accuracy, showing that each submodel performs well with respect to predicting the response variable. When considered together, these two histograms show that leukemia subtype prediction can be done accurately with significantly different sets of genes. In other words, interpreting the coefficients of a given ENET submodel is unreliable, as each ENET submodel does not consistently select the same predictors.

**Table 1.**
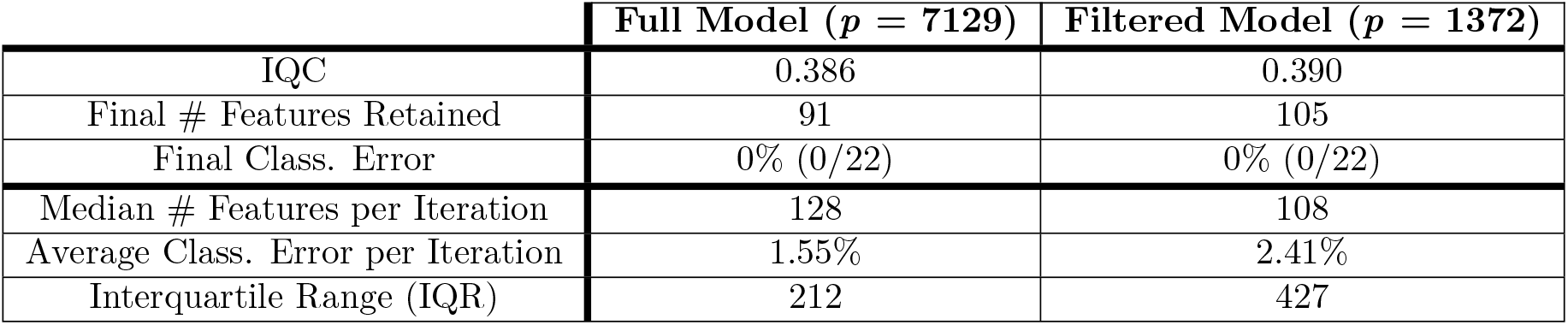
IQC and relevant metrics when applied to Golub et al. microarray data, full and reduced models.

**Fig 4.**
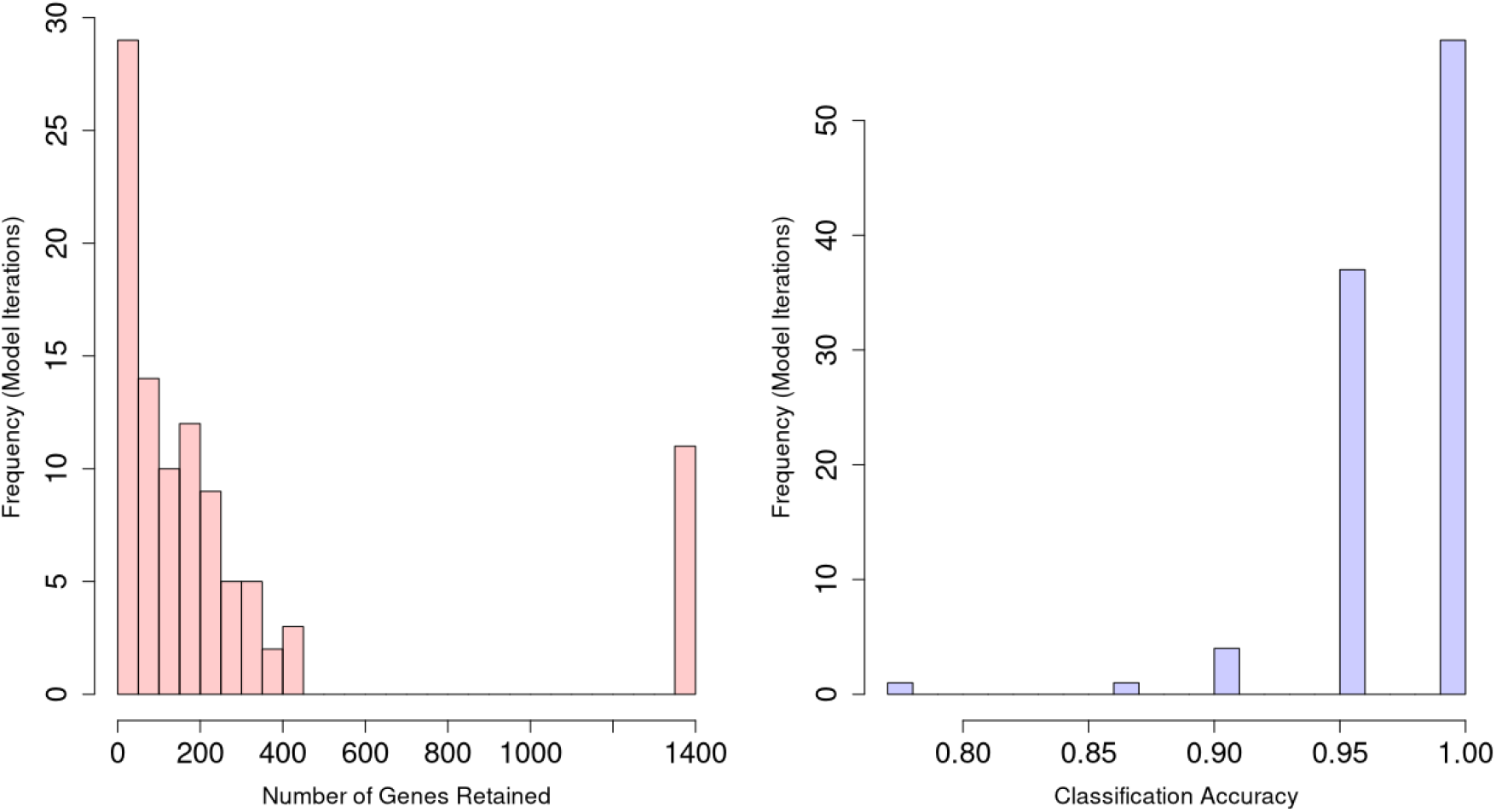
Histograms for number of genes retained and classification accuracy across all 100 microarray ENET submodels. (a) Number of genes retained across all 100 ENET submodels using the filtered data (*n* = 1372). (b) Testing set classification accuracy across all 100 ENET submodels represented as the percentage of individuals given the correct classification of Type I or Type II leukemia.

The original manuscript from [36] selected 50 informative genes as its final predictor set, selecting genes with high correlation to leukemia subtype classification (this was done manually; however, unlike in [8], these 50 genes were analyzed differentially, making for better comparison). We detail differences in the selection of informative genes in Fig 5. To do so, we selected two iterations that had the same number of selected genes, in this case being 54 each. We observed 32 significant genes that were shared among the two models. Of those genes, several aligned with the informative predictors found in the original manuscript, including *Cyclin D3* (S-phase cell cycle progression) and *c-MYB* (oncogene). However, of the genes that differ between the two models, several are cited by [36] to be significant predictors of leukemia subtype, such as *E2A* (oncogene) in Submodel 1, and *Azurocidin* (cationic protein encoder [38]) in Submodel 2. Finally, of the significant predictors cited in the original manuscript, several genes cited to be key predictors of subtype, including *CD11c* (MASPE), *CD33* (mAb surface protein encoder), and *HoxA9* (oncogene), are missing from both iterations.

**Fig 5.**
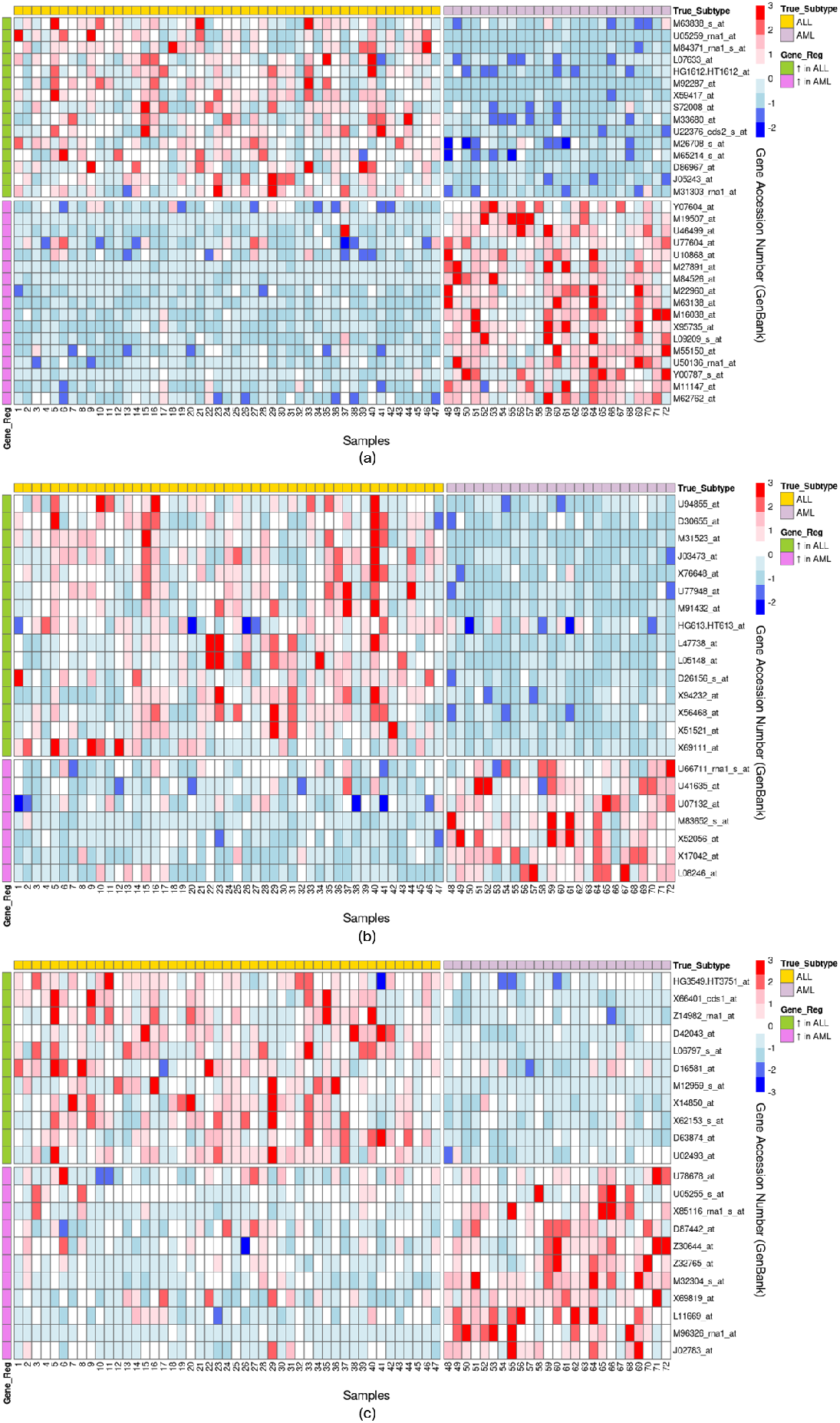
Heatmaps comparing shared and unique genes selected from two IQC-derived submodels (*p* = 54). (a) Genes shared between Submodel 1 and Submodel 2. (b) Genes unique to Submodel 1. (c) Genes unique to Submodel 2. The cells of each heatmap represent normalized expression, centered at 0 with a standard deviation of 1. Expression level minimum/maximum was truncated at ±3 to better visualize difference between groups. Blue cells represent below-mean expression levels, and red-cells represent above-mean expression. Each heatmap is organized by true leukemia subclass (ALL vs. AML), as well as differential expression (genes upregulated in ALL vs. AML).

## Discussion

We have presented IQC as a robust, interpretable metric to quantify and assess the consistency of selected features for ENET. Through our simulation study, we have confirmed via IQC that feature stability in ENET tends to decrease when the data are higher-dimensional, data-poor, and noisy. These results agree with limitations of ENET found in previous literature, especially with respect to data structure [6, 7, 27]. Further, we have shown the capabilities of IQC when applied to real biological data via our microarray case study. Our application illustrates how IQC can be a valuable tool to assess how consistently differentially-expressed genes are preserved across various generations of ENET models. This can then facilitate further investigation of model reliability (see Fig 5). Our application also confirms data modeling issues for biological analyses due to high-dimensionality and low observation counts [21–24].

In our simulation study, we found a steady increase in IQC when observation-to-feature (OTF) ratios are increased in each of the three feature-to-noise (FTN) ratio cases. Interestingly, in both the 25% FTN and 50% FTN simulations, we observed a sharp rise in IQC values at critically low OTF ratios. As we increase the proportion of true features to noisy variables, the behavior of IQC becomes more consistent as OTF decreases. We hypothesize that this behavior is due to ENET becoming increasingly unable to detect any signal within the data. We investigate this in S1 Fig. For a model with *p* = 200 features, we notice that as observation counts decrease, the proportion of features with zero-coefficients increase. This shows that when computing IQC, the comparisons for each individual model are done against essentially a zero-coefficient model, thus resulting in higher “stability” based on our method of computing the metric. However, S1 Fig also shows that an increase of zero-coefficient features also corresponds with an increase in total model error. As such, this behavior is likely due to a very weak signal when training each model iteration given the large number of noisy variables, as well as very few replicates. It is also noteworthy that this behavior is less pronounced in real data, as is the case with our case study. Future work could approach adding weights to the final IQC value based on the magnitude of predictor consistency.

Outside of this oddity, IQC behaves exactly as expected. We anticipated that models with lower dimensionality would provide more stable feature selection, that models with less observations would exhibit higher variance and thus more instability, and that having more noise in the model overall would negatively impact modeling performance based on previous research [6, 7, 27]. Interestingly, while the individual effects of observation counts and feature dimensionality were present, our most significant findings came from looking specifically at the ratio of observations to features. Regardless of the size of the total feature space, models will always tend to be more stable when the ratio of observations to features increases. Importantly, the point at which stability levels out is usually around an OTF of 1.5 across each data configuration. This implies that having more observations than total predictors (significant or otherwise) is a crucial requirement for high stability; however, this is largely unfeasible for most high-dimensional applications such as GWAS and differential expression analysis where the feature space is orders of magnitude larger than the sample size [22–24]. This highlights the importance of understanding model stability when applying methods such as ENET to high-dimensional analyses, and supports our development of IQC.

From our case study, we found that IQC detected higher levels of instability when modeling the full data set compared to the filtered data, reinforcing our simulation results. Additionally, we demonstrated how IQC can be used as a starting point for more in-depth interpretation of ENET coefficient output. Specifically, we highlighted genes cited by [36] as being functionally significant to leukemia classification as being unreliably selected in two models of similar feature space size. Genes such as *CD11c, CD33*, and *MB-1* are particularly useful for distinguishing between ALL and AML lineage cells [36, 39, 40]. However, as noted in the Results section, we found that these particular genes, among others, were not consistently selected in the two highlighted submodels despite known biological significance. This result is important, as it indicates the potential for significant biological insights to be missed or misattributed if using machine learning for *de novo* downstream analyses. Though IQC does not inherently correct the stability issues presented by high-dimensional biological data, it does indicate when further vetting of the model interpretation is necessary. The two selected models each had perfect classification, meaning that each performed well at identifying leukemia subtype. However, any analysis of genes that are differentially expressed will be unreliable due to the inconsistent feature selection. This can indicate the need for corrective procedures such as ensemble modeling that can give coefficient estimates with higher stability [7]. Given the IQC method requires ensemble modeling to derive a comparison model, this aggregate model can also be used as a “final” model for interpretation.

Finally, as shown in the case study, the IQC algorithm is capable of being applied to models where the response variable is non-normal. Though we define IQC through the lens of standard ENET (given its wide application in biological research), IQC can be easily adapted to various other feature selection methods, such as random forest or support vector machines. The sole requirements to generalize IQC are a) modifying the model iteration step to utilize each method’s loss function, and b) redefining how each “optimized” model iteration is selected (Algorithm 2, Step 4; see Methods section). Using our case study as an example, adapting IQC to logistic ENET only involved changing the model function itself (i.e., using logistic ENET model from within glmnet package in R [41]) and redefining model significance to be based on classification error instead of MSE. Thus, we show that IQC is not only robust in the context of ENET, but that it has wider applicability to other modeling infrastructures.

IQC is a novel method for determining model significance via stability, and as such has room for improvements. First, our criterion for generating the aggregate model is somewhat naive. Coefficient retention is determined by a simple majority (i.e., if at least 50% of all model iterations have a non-zero coefficient for feature *j*, that feature is retained in the reference model), and the coefficient values for each nonzero feature are computed by taking the average of each iteration’s coefficient value. While we have shown that IQC is robust and informative, future research can explore more sophisticated aggregation procedures, such as ones proposed by [7]. One such approach could be adding weight to each reference model coefficient *i* based on the proportion of model containing a non-zero coefficient for variable *i*. Additionally, as discussed above, IQC can report “high” stability for models with extremely low observation counts. Future research can explore enhancing the algorithm by incorporating model accuracy weights to account for significantly non-predictive models. This may prove to be easily implemented according to the number of zero-coefficients in a respective reference model, though more testing is necessary. Next, while we presupposed a training/testing split for each model iteration of 70:30 based on convention, we did not explicitly test the impact of this split on IQC. While we selected this ratio to reflect standard practice, previous research cites the benefits of 95:05 and 60:40 splits to accommodate data-limited studies and high overfitting, respectively [31]. Thus, future research exploring less conventional training/testing splits may be warranted. Finally, future usage of IQC may show deviation in the proposed thresholds for low versus high stability. Our proposed thresholds are based on the data available through our simulations, so a more sophisticated thresholding scheme may be explored.

Understanding the consistency of feature selection methods such as ENET is vital for their usage in biological applications. Our work illustrates the robustness of IQC as a means to quantify ENET model stability when applied to health-oriented analyses, and provides a novel approach to integrating model stability evaluation into a more broad analytic infrastructure. Our simulations exhibited an expected, steady increase in IQC when increasing observation counts/OTF ratios, which serves to both confirm the effectiveness of IQC as a model metric, as well as confirm existing knowledge of the effects of high-dimensionality and low sample size on model stability. We then established an appropriate threshold for low, medium, and high stability models based on the distribution of IQC values from our simulations. Our case study illustrated how IQC can be utilized as a means to perform more nuanced *ad hoc* analysis on gene expression data, exploring how standard ENET implementation results in unstable selection of biologically significant genes. Future research can explore tailoring IQC to account for models with low predictive power or mostly zero-coefficients, as well as deeper exploration into a generalized IQC procedure incorporating any generalized loss function or decision rule.

## Materials and methods

### Elastic Net Regression

The IQC method for this article is tailored towards elastic net regression, and thus a short overview of the ENET technique will be provided. (However, as discussed above, the method could be adapted to other similar machine learning techniques.) Elastic net regression is a penalized regression technique which blends two existing regularization methods: lasso, or *l*_1_ penalization, and ridge, or *l*_2_ penalization [8]. Lasso penalties selects for a minimal feature set, while ridge penalties favor many features with small slopes. In short, ENET adds a penalty function to the Ordinary Least Squares (OLS) estimators to mitigate collinearity between predictors and automatically select significant features.

The computation, simplified by [41], is as follows: begin with the response **Y** = (*y*_1_, …, *y*_*n*_), and feature matrix **X** = (*x*_1_ |… |*x*_*k*_), where **x**_*j*_ = (*x*_1*j*_, …, *x*_*nj*_)^*T*^ *j* = 1, …, *p* represents the *j*^*th*^ feature. Standard OLS takes the form **Y** = *β*_0_ + *x*^*T*^ *β* + *ϵ*, where *β*_0_ is the model intercept, *β* are the coefficients for all *p* features, and *ϵ* is the error vector. For ENET, **Y** and **X** are scaled such that **Y** is centered and **X** is standardized as follows:

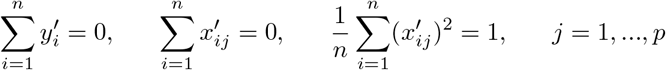

where 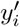 is the centered response for the *i*^*th*^ observation, and 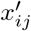 is the standardized measurement for the *i*^*th*^ observation of the *j*^*th*^ feature. Once the data are preprocessed, ENET is computed similarly to OLS such that we minimize model error. Whereas OLS minimizes the sum of squared residuals, ENET minimizes the *penalized* sum of squared residuals, as follows:

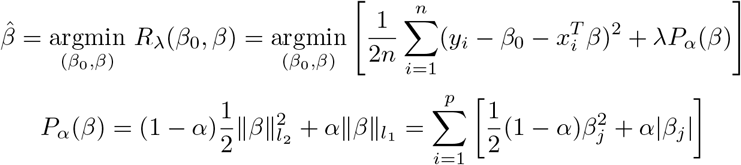

where *α* and *λ* are the hyperparameters for ENET, controlling the mixture and weight of penalization, respectively. When *α* = 1, the model effectively becomes lasso regression, where the penalty function is the *l*_1_ norm of the coefficient vector. Conversely, when *α* = 0, the model becomes ridge regression, and the penalty function is the *l*_2_ norm of the coefficient vector. Finally, *λ* scales the amount of penalization applied to the final coefficient estimates. For a given value of *α*, a grid of *λ* values (typically 100) is generated during the cross validation process to find the optimal hyperparameter combination. The maximum value of this grid, *λ*_*max*_, is defined as the minimum value of *λ* such that all coefficient estimates equal 0 for a given value of *α* [42]. The optimal value for *λ* is then selected such that model error is minimized.

### Data Simulation

The methods for this algorithm require explanation of a few key components. We primarily utilized simulated data to control for the number of features, the number of observations, the error vector, the response variable, our true model coefficients, and which variables are selected to generate the response. Our emphasis with data simulation for this project involved creating a data set that attempts to mimic real-world data. More specifically, we aimed to develop a data set with the following properties:

- Ability to tailor the number of features and observations.
- A collinearity structure that is generalizable; in other words, we are capable of simulating random collinearity between predictors.
- A collinear structure that has natural groupings. As noted in [8], ENET tends to group collinear variables together. We aimed to simulate data in a way that created distinct blocks of collinearity.
- Ability to adjust the distribution of the predictors with relative ease.

With these properties in mind, we developed a robust data simulation method to provide us with controllable, labeled data sets to use as validation for IQC. The data simulation algorithm is detailed in Algorithm 1. From here, we define the algorithm for calculating IQC, which is detailed in Algorithm 2. Our data simulation framework achieves our primary goals with respect to testing IQC. Simulating covariance via sub-covariance block matrices allows for smaller group collinearity and tests of stability against ENET models containing the grouping effect. Finally, generating the response variable from a random sampling of the simulated feature set allows for a) knowledge of true significant predictors, and b) features derived from within and between covariance blocks. The latter point also allows for natural collinearity between true predictors and insignificant predictors. An overview of the data simulation and IQC workflow is provided in Fig 6, and the full derivation of each step is illustrated in Fig 7.

**Fig 6.**
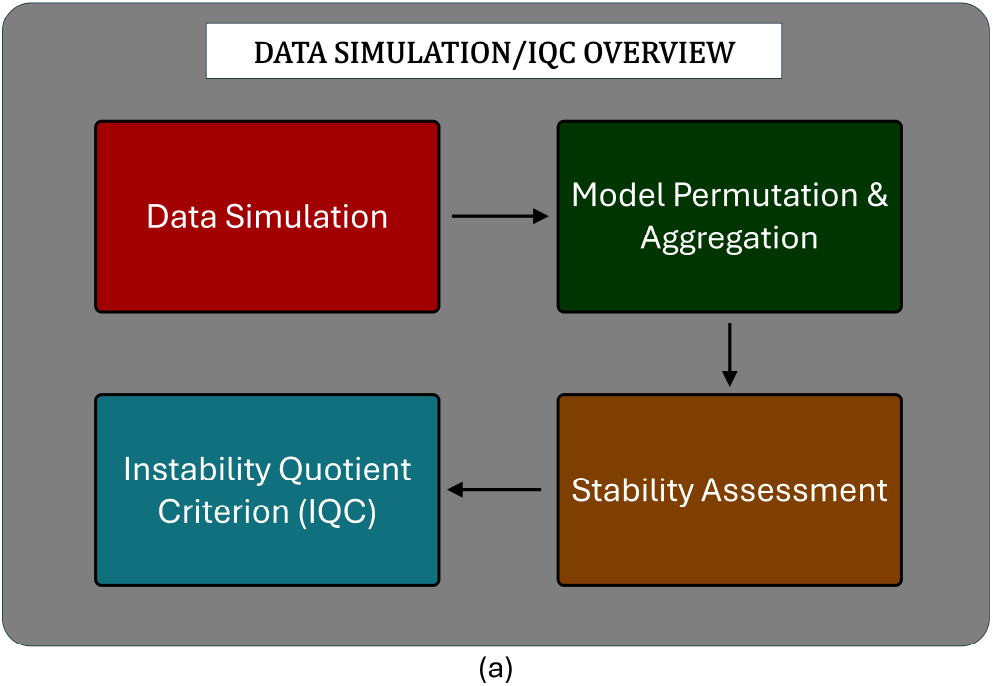
Overview of the workflow for simulating data and conducting IQC. The four major components comprising Algorithms 1 and 2 are Data Simulation, Model Permutation and Aggregation, Stability Assessment, and IQC. This serves as a summarization of the steps taken to develop IQC using our simulated data framework.

**Fig 7.**
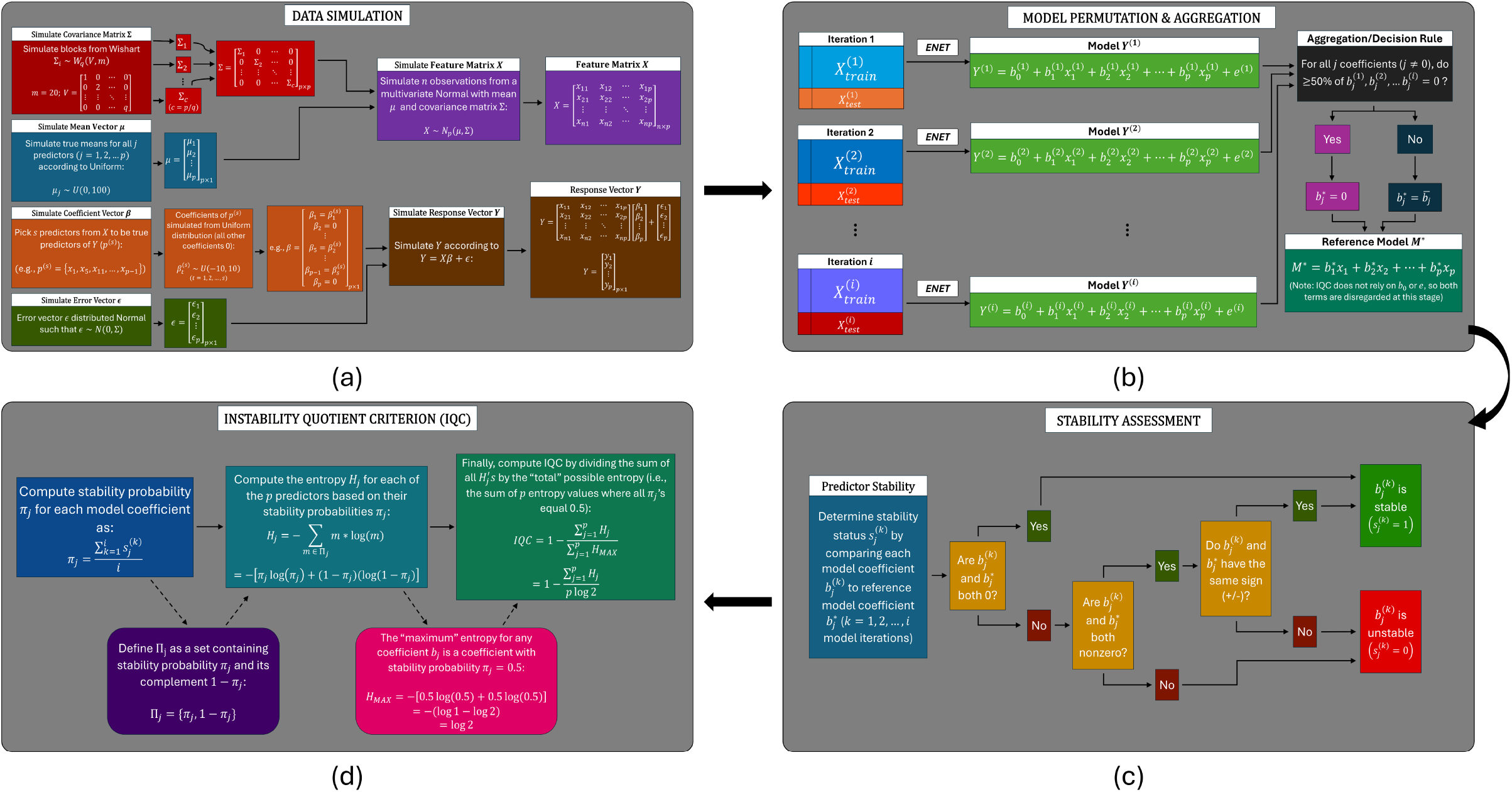
Full derivation of the Data Simulation and IQC workflow. (a) Data Simulation. (b) Model Permutation and Aggregation. (c) Stability Assessment. (d) Instability Quotient Criterion (IQC). The first of the four stages (Data Simulation) represents the steps of Algorithm 1, with the remaining three stages comprising Algorithm 2. Each stage fully details all steps described in Algorithms 1 and 2, along with providing examples of several components of the algorithms (e.g., covariance block sub-matrices Σ_*i*_, example model permutations *Y* ^(*k*)^, etc.).

### Leukemia Microarray Data

Finally, we discuss the methods for preparing the microarray gene expression data from [36]. The raw data contain 7129 gene expression profiles from the bone marrow RNA of 72 subjects with acute leukemia via Affymetrix microarrays. Each individual has a known leukemia subtype diagnosis of either Type I (acute lymphoblastic leukemia, or ALL) or Type II (acute myeloid leukemia, or AML). The response variable *Y* for this data set is a binary representation of each individual’s true leukemia subtype. That is, an individual has a subtype label of 0 if they possess Type I leukemia, and a label of 1 if they possess Type II.

#### Algorithm 1: Method for simulating data used for testing and developing the stability metric

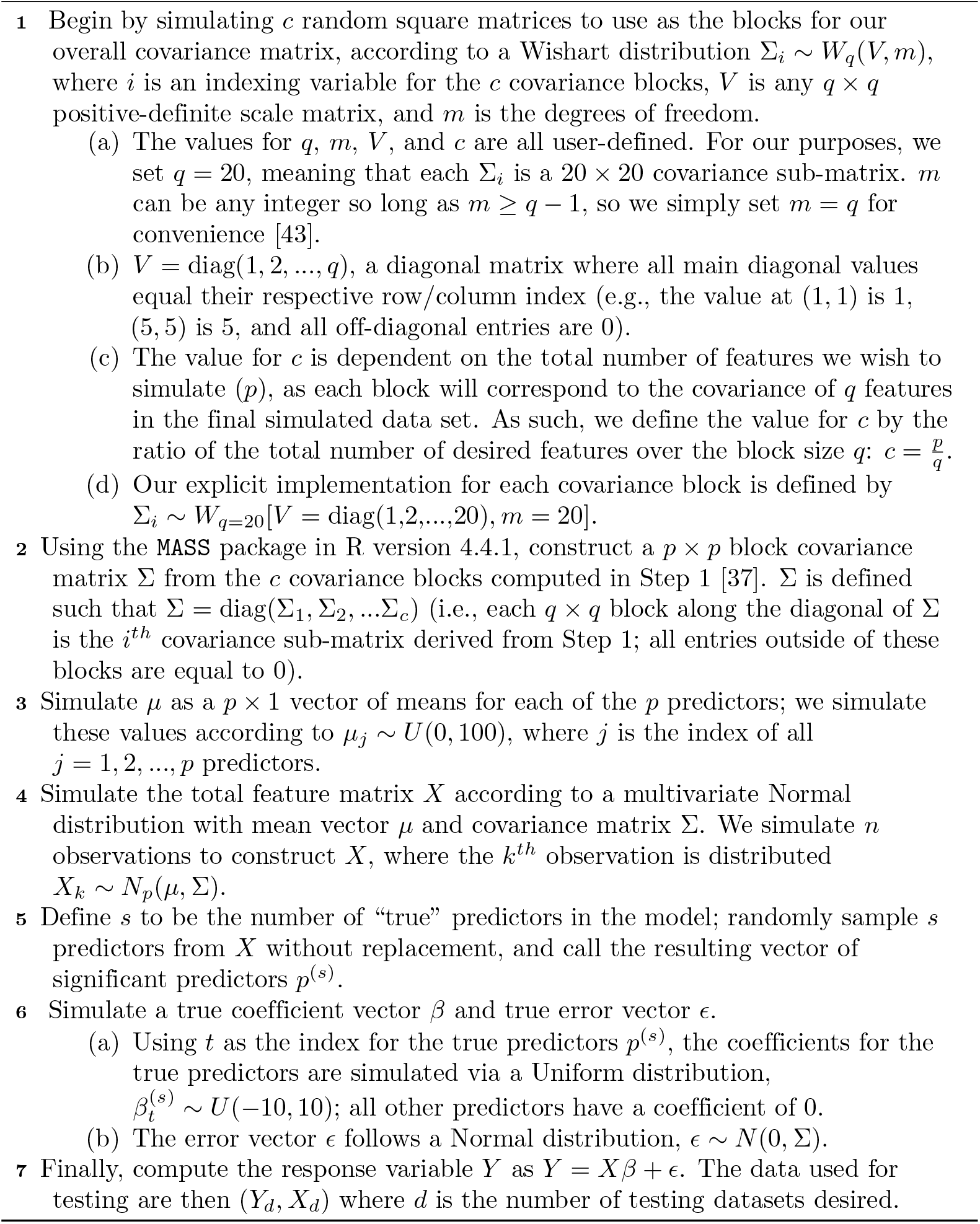

In both the original microarray paper and the original ENET paper, a pre-modeling filtering step was conducted to aid in computational cost [8, 36]. This step was done in both analyses and preserved approximately 1000 genes prior to applying ENET. Because these filtered data sets are not available from their respective publishers, we developed our own filtering process to approximate this step as closely as possible. Pre-filtering was performed by removing reads with a sufficiently low expression interquartile range to reflect existing practices [44]. Genes were removed if their unnormalized interquartile range was less than 1000; this preserved 1372 total genes for feature selection and stability assessment via IQC (S2 Fig displays a histogram of all IQR values for the raw expression data).

#### Algorithm 2: Algorithm for defining Instability Quotient Criterion (IQC)

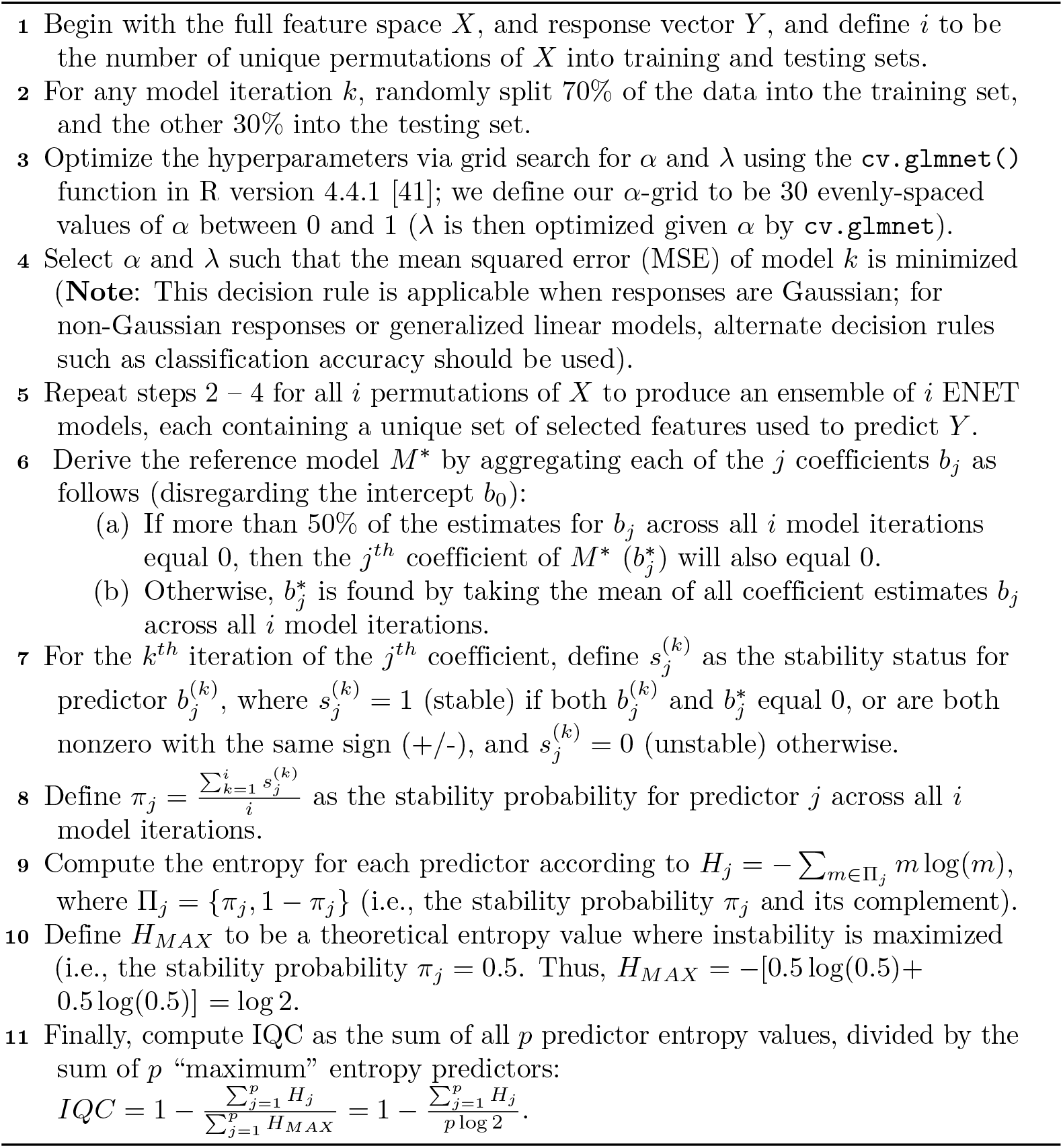

For conducting IQC on the microarray data, we modified Step 4 of Algorithm 2 to utilize classification accuracy for model selection over mean squared error; this was done to account for our binary response variable *Y* representing leukemia subtype. Subsequently, we utilized logistic ENET for model iteration fitting via the glmnet package of R version 4.4.1 [37, 41]. Our predicted response *Ŷ* is then encoded via the indicator function *I*(*ŷ*_*i*_ *>* 0.5) to predict the subtype label for each individual. *y*_*i*_ values greater than 0.5 indicate Type II leukemia, whereas values less than 0.5 indicate Type I.

Finally, when computing IQC, minimum classification error for the predicted response *Ŷ* is used to determine model significance, as opposed to mean squared error for IQC containing standard ENET models. For *ad hoc* analysis of stability, we select two model iterations containing a similar number of significant (retained) genes to the original manuscripts. We utilize the pheatmap package in R version 4.4.1 to visualize differential expression for leukemia subtype, as well as to assess the stability of selecting known, significant genes across multiple model iterations [36, 45].

## Acknowledgments

We would like to thank and Dr. Scott Nuismer, Dr. Terence Soule, and Dr. Tiantian Yang for providing valuable feedback during the development of the data simulation and IQC algorithms.

## References

1. Semmelrock H, Kopeinik S, Theiler D, Ross-Hellauer T, Kowald D. Reproducibility in Machine Learning-Driven Research. arXiv preprint 230710320. 2023.

2. National Academies of Sciences and Medicine and Policy and Global Affairs and Board on Research Data and Division on Engineering and Physical Sciences and Committee on Applied and Theoretical Statistics and Board on Mathematical Sciences and others. Reproducibility and replicability in science. National Academies Press; 2019.

3. Errington TM, Mathur M, Soderberg CK, Denis A, Perfito N, Iorns E, et al. Investigating the replicability of preclinical cancer biology. elife. 2021;10:e71601.

4. Nuijten MB. Practical tools and strategies for researchers to increase replicability. Developmental Medicine & Child Neurology. 2019;61(5):535–9.

5. Riley RD, Collins GS. Stability of clinical prediction models developed using statistical or machine learning methods. Biometrical Journal. 2023;65(8):2200302.

6. Wang W, Liang J, Liu R, Song Y, Zhang M. A Robust Variable Selection Method for Sparse Online Regression via the Elastic Net Penalty. Mathematics. 2022;10(16). Available from: https://www.mdpi.com/2227-7390/10/16/2985. xdoi:10.3390/math10162985.

7. Jenul A, Schrunner S, Liland KH, Indahl UG, Futsæther CM, Tomic O. Rent—repeated elastic net technique for feature selection. IEEE Access. 2021;9:152333–46.

8. Zou H, Hastie T. Regularization and variable selection via the elastic net. Journal of the Royal Statistical Society: Series B (Statistical Methodology). 2005;67(2): 301–20.

9. Binka M, Klaver B, Cua G, Wong AW, Fibke C, Velásquez García HA, et al. An Elastic Net Regression Model for Identifying Long COVID Patients Using Health Administrative Data: A Population-Based Study. In: Open Forum Infectious Diseases. vol. 9. Oxford University Press US; 2022. ofac640.

10. Liu W, Li Q. An efficient elastic net with regression coefficients method for variable selection of spectrum data. PloS one. 2017;12(2):e0171122.

11. Moxley TA, Johnson-Leung J, Seamon E, Williams C, Ridenhour BJ. Application of elastic net regression for modeling COVID-19 sociodemographic risk factors. Plos one. 2024;19(1):e0297065.

12. Xie J, Cho H, Lin BM, Pillai M, Heimisdottir LH, Bandyopadhyay D, et al. Improved metabolite prediction using microbiome data-based elastic net models. Frontiers in Cellular and Infection Microbiology. 2021:972.

13. Chakraborty S, Tomsett R, Raghavendra R, Harborne D, Alzantot M, Cerutti F, et al. Interpretability of deep learning models: A survey of results. In: 2017 IEEE smartworld, ubiquitous intelligence & computing, advanced & trusted computed, scalable computing & communications, cloud & big data computing, Internet of people and smart city innovation (smartworld/SCALCOM/UIC/ATC/CBDcom/IOP/SCI). IEEE; 2017. p. 1–6.

14. Tibshirani R. Regression shrinkage and selection via the lasso. Journal of the Royal Statistical Society Series B: Statistical Methodology. 1996;58(1):267–88.

15. Vaarhorst A. Genetic and metabolomic approaches for coronary heart disease risk prediction. European heart journal. 2003;24(11):987–1003.

16. Hoerl AE, Kennard RW. Ridge regression: Biased estimation for nonorthogonal problems. Technometrics. 1970;12(1):55–67.

17. Hoerl RW. Ridge regression: a historical context. Technometrics. 2020;62(4):420–5.

18. Marquardt DW, Snee RD. Ridge regression in practice. The American Statistician. 1975;29(1):3–20.

19. Neufeld A, Dharamshi A, Gao LL, Witten D. Data thinning for convolution-closed distributions. Journal of Machine Learning Research. 2024;25(57):1–35.

20. Albani V, Loria J, Massad E, Zubelli J. COVID-19 underreporting and its impact on vaccination strategies. BMC Infectious Diseases. 2021;21:1–13.

21. Mokhtari EB, Ridenhour BJ. Filtering ASVs/OTUs via mutual information-based microbiome network analysis. BMC bioinformatics. 2022;23(1):380.

22. Chen R, Yang L, Goodison S, Sun Y. Deep-learning approach to identifying cancer subtypes using high-dimensional genomic data. Bioinformatics. 2020;36(5):1476–83.

23. Sen PK. Robust statistical inference for high-dimensional data models with application to genomics. Austrian journal of statistics. 2006;35(2&3):197–214.

24. Clarke R, Ressom HW, Wang A, Xuan J, Liu MC, Gehan EA, et al. The properties of high-dimensional data spaces: implications for exploring gene and protein expression data. Nature reviews cancer. 2008;8(1):37–49.

25. Fan J, Lv J. Sure independence screening for ultrahigh dimensional feature space. Journal of the Royal Statistical Society Series B: Statistical Methodology. 2008;70(5):849–911.

26. Gupta S, Gupta A. Dealing with noise problem in machine learning data-sets: A systematic review. Procedia Computer Science. 2019;161:466–74.

27. Zou H, Zhang HH. On the adaptive elastic-net with a diverging number of parameters. Annals of statistics. 2009;37(4):1733.

28. Pereira MB, Wallroth M, Jonsson V, Kristiansson E. Comparison of normalization methods for the analysis of metagenomic gene abundance data. BMC genomics. 2018;19:1–17.

29. Joseph VR. Optimal ratio for data splitting. Statistical Analysis and Data Mining: The ASA Data Science Journal. 2022;15(4):531–8.

30. Muraina I. Ideal dataset splitting ratios in machine learning algorithms: general concerns for data scientists and data analysts. In: 7th international Mardin Artuklu scientific research conference; 2022. p. 496–504.

31. Sivakumar M, Parthasarathy S, Padmapriya T. Trade-off between training and testing ratio in machine learning for medical image processing. PeerJ Computer Science. 2024;10:e2245.

32. Büyükkeçeci M, Okur MC. A comprehensive review of feature selection and feature selection stability in machine learning. Gazi University Journal of Science. 2023;36(4):1506–20.

33. Turkiewicz A, Luta G, Hughes H, Ranstam J. Statistical mistakes and how to avoid them–lessons learned from the reproducibility crisis. Osteoarthritis and Cartilage. 2018;26(11):1409–11.

34. Khan MHR, Bhadra A, Howlader T. Stability selection for lasso, ridge and elastic net implemented with AFT models. Statistical applications in genetics and molecular biology. 2019;18(5).

35. Shannon CE. A mathematical theory of communication. ACM SIGMOBILE mobile computing and communications review. 2001;5(1):3–55.

36. Golub TR, Slonim DK, Tamayo P, Huard C, Gaasenbeek M, Mesirov JP, et al. Molecular classification of cancer: class discovery and class prediction by gene expression monitoring. science. 1999;286(5439):531–7.

37. R Core Team. R: A Language and Environment for Statistical Computing. Vienna, Austria; 2022. Available from: https://www.R-project.org/.

38. Aries PM, Csernok E, Gross WL. Antineutrophil cytoplasmatic autoantibodies with specificity to proteinase 3. In: Autoantibodies. Elsevier; 2007. p. 119–26.

39. Dinndorf PA, Buckley JD, Nesbit ME, Lampkin BC, Piomelli S, Feig SA, et al. Expression of myeloid differentiation antigens in acute nonlymphocytic leukemia: Increased concentration of CD33 antigen predicts poor outcome—A report from the childrens cancer study group. Medical and pediatric oncology. 1992;20(3):192–200.

40. Sakhinia E, Farahangpour M, Tholouli E, Yin JL, Hoyland JA, Byers RJ. Comparison of gene-expression profiles in parallel bone marrow and peripheral blood samples in acute myeloid leukaemia by real-time polymerase chain reaction. Journal of clinical pathology. 2006;59(10):1059–65.

41. Friedman J, Hastie T, Tibshirani R. Regularization Paths for Generalized Linear Models via Coordinate Descent. Journal of Statistical Software. 2010;33(1):1–22. doi:10.18637/jss.v033.i01.

42. Hastie T, Qian J, Tay K. An introduction to glmnet. CRAN R Repositary. 2021;5:1–35.

43. Uhlig H. On singular Wishart and singular multivariate beta distributions. The Annals of Statistics. 1994:395–405.

44. Chockalingam S, Aluru M, Aluru S. Microarray data processing techniques for genome-scale network inference from large public repositories. Microarrays. 2016;5(3):23.

45. Kolde R. pheatmap: Pretty Heatmaps; 2025. R package version 1.0.13. Available from: https://github.com/raivokolde/pheatmap.

