## Supporting information for "*IQC:* A Novel Criterion for Assessing Feature Selection Stability in High-Dimensional Analyses"

### Supporting information 551

**S1 Fig. Comparing the proportion of zero-coefficient features and normalized RMSE for when decreasing observations for ENET ($p = 200$).** 552 553

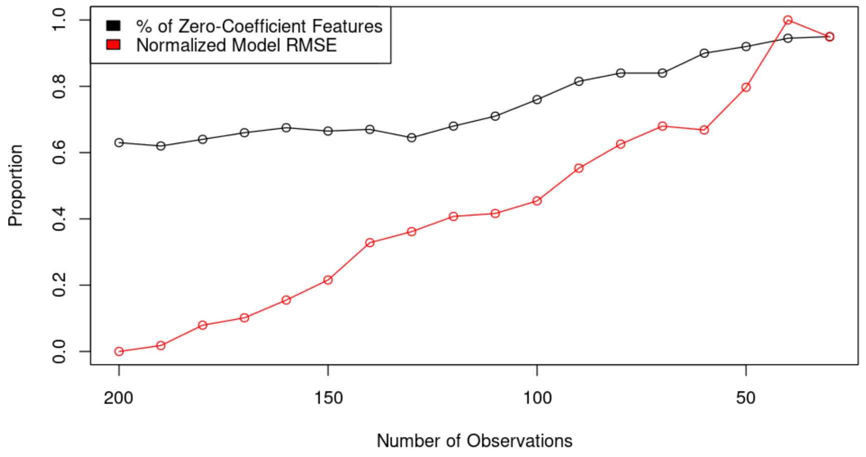


**S2 Fig. Interquartile range for 7129 gene expression profiles of acute leukemia subjects from microarray case study.** 554 555

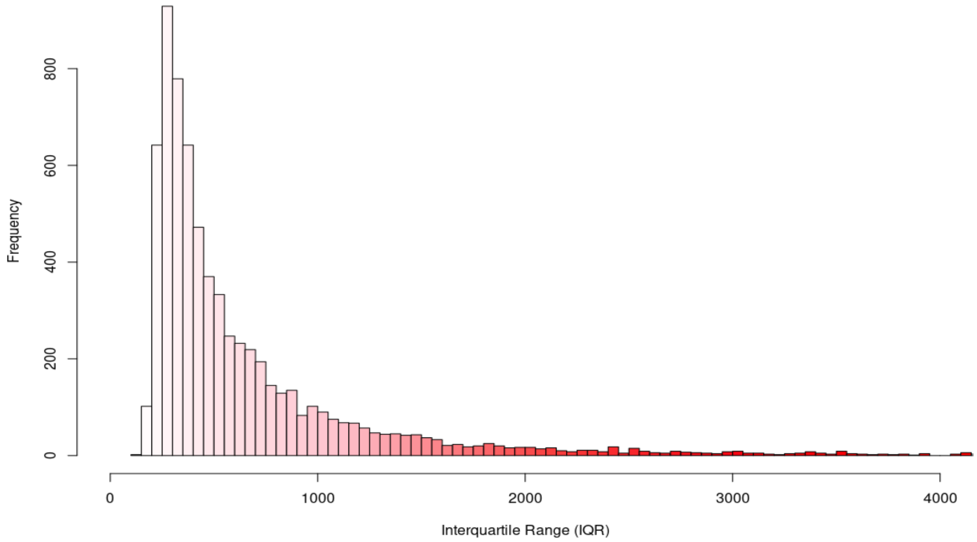

**S1 Table. Complete list of IQC values for each feature-to-noise ratio and number of features, with decreasing observation counts.**

| | # Features 25% FTN Ratio | | | | # Features 50% FTN Ratio | | | | # Features 75% FTN Ratio | | | |
|---|---|---|---|---|---|---|---|---|---|---|---|---|
| **Observations** | **60** | **100** | **140** | **200** | **60** | **100** | **140** | **200** | **60** | **100** | **140** | **200** |
| 200 | 0.797 | 0.732 | 0.692 | 0.477 | 0.815 | 0.713 | 0.592 | 0.418 | 0.853 | 0.790 | 0.565 | 0.395 |
| 190 | 0.755 | 0.730 | 0.678 | 0.459 | 0.805 | 0.704 | 0.572 | 0.400 | 0.839 | 0.775 | 0.536 | 0.359 |
| 180 | 0.731 | 0.713 | 0.659 | 0.438 | 0.798 | 0.682 | 0.540 | 0.406 | 0.842 | 0.765 | 0.495 | 0.342 |
| 170 | 0.731 | 0.693 | 0.655 | 0.399 | 0.804 | 0.648 | 0.505 | 0.382 | 0.831 | 0.757 | 0.464 | 0.311 |
| 160 | 0.747 | 0.690 | 0.645 | 0.385 | 0.803 | 0.620 | 0.466 | 0.359 | 0.829 | 0.744 | 0.449 | 0.297 |
| 150 | 0.742 | 0.676 | 0.614 | 0.374 | 0.774 | 0.600 | 0.443 | 0.351 | 0.820 | 0.610 | 0.428 | 0.314 |
| 140 | 0.718 | 0.645 | 0.574 | 0.346 | 0.769 | 0.580 | 0.415 | 0.327 | 0.805 | 0.565 | 0.424 | 0.317 |
| 130 | 0.691 | 0.629 | 0.542 | 0.339 | 0.749 | 0.544 | 0.375 | 0.313 | 0.790 | 0.514 | 0.423 | 0.301 |
| 120 | 0.632 | 0.602 | 0.503 | 0.334 | 0.713 | 0.505 | 0.346 | 0.311 | 0.796 | 0.489 | 0.408 | 0.302 |
| 110 | 0.670 | 0.562 | 0.469 | 0.340 | 0.717 | 0.470 | 0.329 | 0.291 | 0.772 | 0.451 | 0.371 | 0.296 |
| 100 | 0.653 | 0.535 | 0.432 | 0.362 | 0.684 | 0.437 | 0.311 | 0.307 | 0.761 | 0.431 | 0.346 | 0.267 |
| 90 | 0.647 | 0.472 | 0.370 | 0.371 | 0.571 | 0.327 | 0.304 | 0.254 | 0.518 | 0.370 | 0.305 | 0.236 |
| 80 | 0.610 | 0.409 | 0.366 | 0.383 | 0.485 | 0.299 | 0.287 | 0.249 | 0.494 | 0.340 | 0.300 | 0.248 |
| 70 | 0.551 | 0.321 | 0.386 | 0.385 | 0.424 | 0.260 | 0.328 | 0.242 | 0.455 | 0.291 | 0.267 | 0.264 |
| 60 | 0.469 | 0.302 | 0.366 | 0.455 | 0.365 | 0.282 | 0.451 | 0.338 | 0.361 | 0.237 | 0.374 | 0.346 |
| 50 | 0.383 | 0.244 | 0.400 | 0.305 | 0.325 | 0.218 | 0.329 | 0.348 | 0.306 | 0.254 | 0.493 | 0.444 |
| 40 | 0.363 | 0.166 | 0.427 | 0.743 | 0.289 | 0.311 | 0.393 | 0.296 | 0.218 | 0.271 | 0.482 | 0.293 |
| 30 | 0.253 | 0.481 | 0.399 | 0.535 | 0.292 | 0.349 | 0.181 | 0.692 | 0.479 | 0.403 | 0.517 | 0.337 |
